# The G protein-Coupled Metabotropic Glutamate Receptor 1 controls neuronal macroautophagy

**DOI:** 10.1101/2020.11.02.365783

**Authors:** Maribel Donoso, Luisa Speranza, Magdalena Kalinowska, Catherine Castillo, Claudia De Sanctis, Anna Francesconi

## Abstract

Autophagy is an evolutionarily conserved, highly regulated catabolic process critical to neuronal homeostasis, function and survival throughout organismal lifespan. However, the external factors and signals that control autophagy in neurons are still poorly understood. Here we report that the G protein-coupled metabotropic glutamate receptor 1 (mGlu1) contributes to control basal autophagy in the brain. Autophagy is upregulated in the brain of adult mGlu1 knockout mice and genetic deletion or pharmacological inhibition of native mGlu1 receptors enhances autophagy flux in neurons. The evolutionarily conserved adaptor protein FEZ1, identified by a genome-wide screen as mGlu1 receptor interacting partner, was found to participate in the regulation of neuronal autophagy and to be required for repression of autophagy flux by the mGlu1 receptor. Furthermore, FEZ1 appears to enable association of mGlu1 with Ulk1, a core component of the autophagy pathway. Thus, we propose that the mGlu1 receptor contributes to restrain constitutive autophagy in neurons.

## Introduction

Rare missense and frame-shift mutations in the *GRM1* gene encoding metabotropic glutamate receptor 1 (mGlu1) result in congenital cerebellar ataxia, global developmental delays and moderate to severe intellectual deficit (Davarniya et al., 2015; Guergueltcheva et al., 2012) (Watson et al., 2017). Brain imaging studies of affected individuals also describe progressive involution of the cerebellum and a constitutionally small brain in some subjects (Guergueltcheva et al., 2012). MGlu1 is an evolutionarily conserved GPCR belonging to the mGluR subfamily of eight CNS-expressed receptors (mGlu1 through mGlu8) that modulate synaptic transmission and neuronal excitation (Niswender and Conn, 2010; Reiner and Levitz, 2018). In both human and murine genome the mGlu1 receptor comprises four alternatively spliced variants - differing in the composition of the carboxyl terminal tail - that are broadly expressed in the brain and chiefly present in neurons (Berthele et al., 1998; Naito et al., 2018). Mutations in patients give rise to nonfunctional receptors by either impairing glutamate binding or by generating transcripts that lack the 7-transmembrane region or encode abnormal cytoplasmic tails. Global deletion of *Grm1* in mice (*Grm1*^−/−^) recapitulates pathological manifestations including ataxia and disturbances in learning and memory as indicated by deficits in hippocampal LTD, LTP and associative learning (Aiba et al., 1994a; Aiba et al., 1994b; Conquet et al., 1994; Volk et al., 2006). The pathological manifestations of mGlu1 silencing can be explained by its critical role in the establishment and maintenance of neural circuits (Kano et al., 1997; Narushima et al., 2016; Narushima et al., 2019) and formation and remodeling of dendritic spines (Kalinowska et al., 2015; Sugawara et al., 2017), the site of excitatory synapses. However, the profound deficits associated with loss of mGlu1 activity are not adequately explained by current knowledge of its signaling and cellular functions. The mGlu1 receptor couples *via* Gα_q/11_ to phospholipase C leading to production of InsP3 which mobilizes Ca^2+^ from the endoplasmic reticulum. Stimulation of the mGlu1 receptor also activates extracellular signal-regulated kinase/ERK (Karim et al., 2001; Kumari et al., 2013) and mechanistic target of rapamycin/mTOR kinase (Hou and Klann, 2004). mTOR is a nutrient sensor that controls cellular anabolism and catabolism through its capacity to promote protein synthesis and suppress macroautophagy, respectively (Kim et al., 2002; Kim and Guan, 2019; Liu and Sabatini, 2020; Saxton and Sabatini, 2017). Accordingly, the capacity of the mGlu1 receptor to stimulate protein synthesis in neurons was documented (Yu et al., 2013) but whether it can influence neuronal catabolism *via* macroautophagy is unknown.

Macroautophagy (autophagy hereafter) is a highly regulated catabolic process whereby cytoplasmic content is encapsulated within double membrane vesicles, termed autophagosomes, through which the cargo is delivered to lysosomes for degradation. The underpinnings of this process involve a group of evolutionarily conserved autophagy-related *(ATG)* genes that encode proteins necessary for autophagosome formation, expansion and fusion with lysosomes (Mizushima et al., 2011; Xie and Klionsky, 2007). Autophagy maintains cell and tissue homeostasis by regenerating nutrients in response to environmental stress such as starvation (Morishita and Mizushima, 2019). An additional critical housekeeping function is the selective removal and digestion of protein aggregates and damaged organelles such as mitochondria and peroxisomes (Anding and Baehrecke, 2017; Gatica et al., 2018; Lim and Yue, 2015) that are engulfed in autophagosomes through recognition by autophagy cargo receptors (Kirkin et al., 2009; Pohl and Dikic, 2019; Zaffagnini and Martens, 2016).

The maturation (Ban et al., 2013; Clark et al., 2018; Shen and Ganetzky, 2009; Stavoe et al., 2016; Stavoe and Holzbaur, 2019) and survival (Hara et al., 2006; Komatsu et al., 2006; Komatsu et al., 2007) of neurons is dependent on autophagy and its dysfunction is linked to neurodegenerative conditions including Huntington’s and Alzheimer’s disease and spinocerebellar ataxia (SCA3) among others (Menzies et al., 2017; Nixon, 2013). In vertebrates, autophagy is basally active in most cells and tissues and strongly upregulated by starvation (Kuma et al., 2004; Mizushima et al., 2004). However, starvation has little effect in inducing autophagy in the brain and neurons (Kaushik et al., 2011; Mizushima et al., 2004; Nikoletopoulou et al., 2017) in which basal autophagy is fairly efficient (Boland et al., 2008; Maday and Holzbaur, 2014). Neurons are post-mitotic cells the lifespan of which, in absence of disease, parallels the duration of organismal lifespan (Magrassi et al., 2013): as such, they need sensitive and reliable mechanisms to gauge their metabolic state to maintain metabolic and structural homeostasis. Moreover, in the CNS efficient autophagy needs to be maintained and fine-tuned past initial phases of developmental growth, to support neuronal health and structural and functional homeostasis of established circuits in the adult (Lieberman et al., 2020; Stavoe and Holzbaur, 2020). Despite the progress in understanding the mechanisms of autophagy and its critical role in the CNS, we still have limited knowledge of the external cues and signals that control the initiation and progression of neuronal autophagy in physiological conditions (Nikoletopoulou and Tavernarakis, 2018). The potential role of neurotransmitter receptors in the regulation of autophagy is beginning to receive attention (Shehata et al., 2012; Yue et al., 2002) but is as yet little understood.

Here, we report that the mGlu1 receptor represses constitutive neuronal autophagy. We found that constitutive autophagy is enhanced in the brain of *Grm1*^−/−^ mice and that genetic deletion of mGlu1 or pharmacological inhibition of its activity enhances autophagy flux in neurons, whereas transient receptor activation represses autophagy. Mechanistically, we found that mGlu1 interacts with the evolutionary conserved adaptor protein FEZ1, which we show to be required for efficient neuronal autophagy. Interaction with FEZ1 is necessary for mGlu1 receptor capacity to repress autophagy flux in neurons and mediates mGlu1 association with core components of the autophagy pathway. Together these findings converge to support a critical function of mGlu1 receptors in restraining constitutive autophagy in the CNS.

## Results

### Loss of mGlu1 receptor causes an imbalance of basal autophagy in the brain

To begin examining whether mGlu1 receptor activity could contribute to the regulation of neuronal catabolism, we surveyed autophagy in the brain cortex of mature adult mGlu1 knockout mice (*Grm1*^−/−^ mice) that lack all mGlu1 receptor variants (Conquet et al., 1994). The mGlu1 receptor is broadly distributed in the cortex with peak expression in the adult, in both rodent and human brain (Boer et al., 2010; Ong et al., 1998; Shigemoto et al., 1992). *Grm1*^−/−^ mice display no overt phenotype until the second to third postnatal week, when they develop ataxia and intention tremor that however do not compromise longevity. To assess autophagy, we first examined the phosphatidylethanolamine-conjugated form of microtubule-associated protein 1 light chain 3b (LC3b) - a selective marker of autophagosomes (Klionsky et al., 2016). When autophagy is induced, cytosolic LC3 (LC3-I) is converted by lipidation to LC3-II, which associates with autophagosome membranes. Thus the amount of LC3-II at steady state can provide a proxy measure of LC3-I conversion. Using immunoblot, we found that LC3-II was more abundant in cortical extracts of *Grm1*^−/−^ mice compared to *Grm1^+/+^* littermates *(**Figure 1A***). To examine autophagy *in situ*, we used anti-LC3b and fluorescent labeling of brain sections to visualize LC3b^+^ autophagic vacuoles (Klionsky et al., 2016) (but see (Runwal et al., 2019)). Labeled LC3b exhibits both diffuse and punctate distribution arising from its association with autophagic vacuoles (Klionsky et al., 2016). In cortical regions, LC3b^+^ puncta were clearly discernible in neuronal soma and more abundant in *Grm1*^−/−^ mice compared to wild type *(**Figure 1B-C***). Together, the increase in LC3-II and number of autophagosomes hinted at perturbations of autophagy in *Grm1*^−/−^ mice but do not distinguish between potential upregulation of autophagosome formation or block of autophagic degradation.

**Figure 1.**
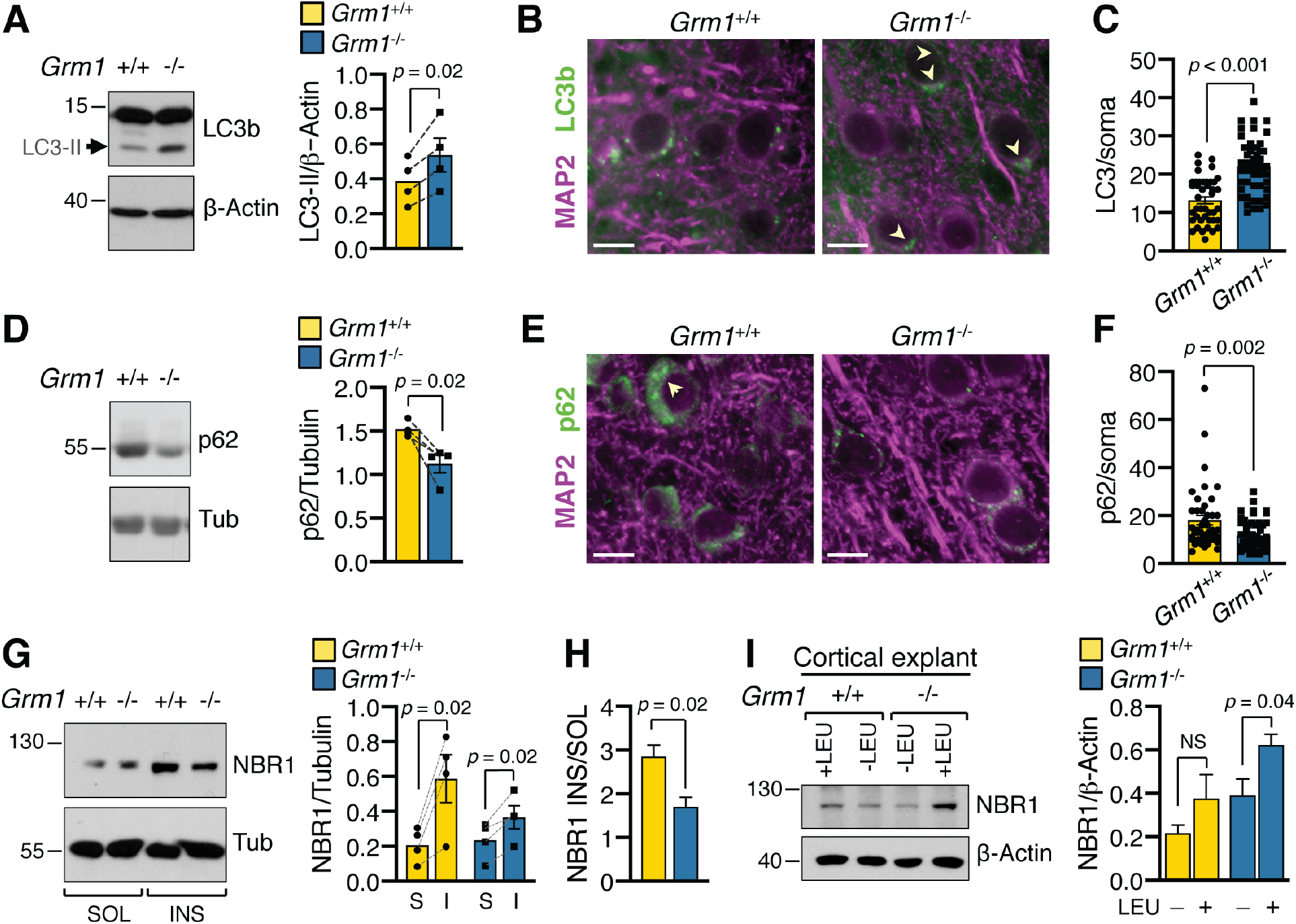
Mice lacking mGlu1 receptor show imbalance of autophagy in the brain. (**A**) Immunoblot and quantification of LC3-II normalized to β-actin (mean±SEM). Symbols in bar graphs represent individual matched littermates (6-8 month-old); N=4 mice per group, paired t-test. (**B**) Representative images of cortical fields from brain sections labeled with anti-LC3b and anti-MAP2 to visualize the somatodendritic compartment, scale bars 25 μm. Arrowheads point to clusters of LC3^+^ puncta. (**C**) Quantification of LC3^+^ puncta in neuronal soma from images like those in (B). Symbols represent LC3^+^ counts in individual neurons from matched littermates: *Grm1^+/+^* n=42 neurons, *Grm1*^−/−^ n=53, N=4 mice (12 month-old) per group, Mann-Whitney test. (**D**) Immunoblot and quantification of p62 normalized to tubulin (mean±SEM): N=4 matched littermates mice per group, paired t-test. (**E**) Representative images of cortical fields from brain sections labeled with anti-p62 and anti-MAP2, scale bars 25 μm. (**F**) Quantification of p62 in neuronal soma from images like those in (E). p62^+^ puncta per cell *Grm1^+/+^* n=44 neurons, *Grm1*^−/−^ n=53 from N=4 mice per group, Mann-Whitney test. (**G**) Immunoblot and quantification of soluble and insoluble NBR1 in brain cortex. NBR1 normalized to tubulin (mean±SEM), N=4 mice, paired t-test. (**H**) NBR1 enrichment in insoluble fraction (INS/SOL; mean±SEM), N=4 mice per group, unpaired t-test. (**I**) Immunoblot and quantification of NBR1 in tissue explants incubated with and without leupeptin (LEU). Mean±SEM, *Grm1^+/+^*-LEU n=6, +LEU n=5, *p*=0.173; *Grm1*^−/−^ −LEU n=6, +LEU n=5, *p*=0.04, unpaired t-test; n, independent determinations from 3 mice per group.

Sequestosome 1/p62 (p62 hereafter) is an ubiquitous autophagy receptor that participates in the removal of ubiquitinated targets (Bjorkoy et al., 2009; Kirkin et al., 2009; Lamark et al., 2017; Pankiv et al., 2007). p62 is recruited to autophagosomes and degraded together with their cargo so that its relative abundance is inversely correlated to autophagy flux. At steady state, both p62 expression determined by immunoblot *(**Figure 1D**)*, and p62^+^ puncta in neuronal soma detected by immunolabeling *(**Figure 1E-F**)* were decreased in cortical tissue of *Grm1*^−/−^ mice compared to wild type littermates. We considered the possibility that differences in the expression of LC3b and p62 could arise from changes in their rate of transcription that has been documented (Fullgrabe et al., 2016; Settembre et al., 2011). However mRNA expression of *Map1lc3b* and *Sqstm1*, encoding LC3b and p62 respectively, was comparable between genotypes as determined by RT-PCR *(**Figure 1 – figure supplement 1**)* suggesting that differences in gene expression are unlikely to underlie the observed phenotypes.

In addition to p62, other autophagy cargo receptors are expressed in the CNS including neighbor of BRCA1 gene 1 (NBR1) which is also degraded by autophagy independently of p62 (Kirkin et al., 2009). NBR1 was shown to co-fractionate predominantly with insoluble material in adult human brain, likely due to association with endogenous protein aggregates (Odagiri et al., 2012). Thus we used immunoblot to examine NBR1 expression in soluble and insoluble fractions extracted from the brain cortices of adult mice. Although partly soluble, NBR1 was appreciably enriched in the insoluble fraction in wild type mice: in contrast, in *Grm1*^−/−^ mice the relative abundance of insoluble NBR1 was significantly reduced *(**Figure 1G-H)**.* NBR1 was shown to undergo more rapid lysosomal turnover than p62, at least in human HeLa cells (Kirkin et al., 2009). We hypothesized that the reduction in insoluble NBR1 at steady state could be indicative of enhanced rate of clearance *via* autophagy. To examine this possibility, we performed *ex vivo* autophagy flux assays by incubating (60 min) freshly microdissected cortical tissue explants with leupeptin, an inhibitor of serine and cysteine proteases, together with ammonium chloride and examined NBR1 abundance by immunoblot. In the presence of the lysosomal inhibitors, undigested NBR1 was modestly increased in wild-type explants; in contrast, incubation with the inhibitors resulted in robust NBR1 accumulation in tissue from *Grm1*^−/−^ mice *(**Figure 1I**)*, an effect that could arise from increased rate of autophagy flux. Thus, taken together these findings suggest that loss of mGlu1 receptor results in upregulation of autophagy *in vivo.*

### MGlu1 receptor activity represses basal autophagy in neurons

Autophagy is a highly dynamic process and homeostatic alterations observed in brain tissue may not reflect perturbations in autophagy flow directly dependent on mGlu1 receptor activity. Thus we examined autophagy flux in dissociated hippocampal neurons from *Grm1*^−/−^ and wild type mouse pups. For this, neurons maintained *in vitro* under basal conditions replete with nutrients were incubated in the absence or presence of bafilomycin A_1_, (bafA_1_; 30 min), a lysosomal V-ATPase inhibitor that blocks autophagosome–lysosome fusion causing accumulation of autophagy cargo (Yamamoto et al., 1998). We used immunolabeling to visualize the autophagy receptor p62 - which is degraded together with autophagosome cargo upon fusion with lysosomes - focusing on the somatodendritic region where the mGlu1 receptor is enriched (Kalinowska et al., 2015). In wild type neurons, p62^+^ puncta were prominent in the soma and sparse in dendrites and their abundance was marginally enhanced by incubation with bafA1, indicating slow clearance in the presence of nutrients *(**Figure 2A-B***). In contrast, in *Grm1*^−/−^ neurons incubation with bafA_1_ resulted in robust increase of p62^+^ puncta compared to untreated *(**Figure 2A-B**)*, indicating a higher rate of p62 degradation *via* autophagy in absence of the mGlu1 receptor.

**Figure 2.**
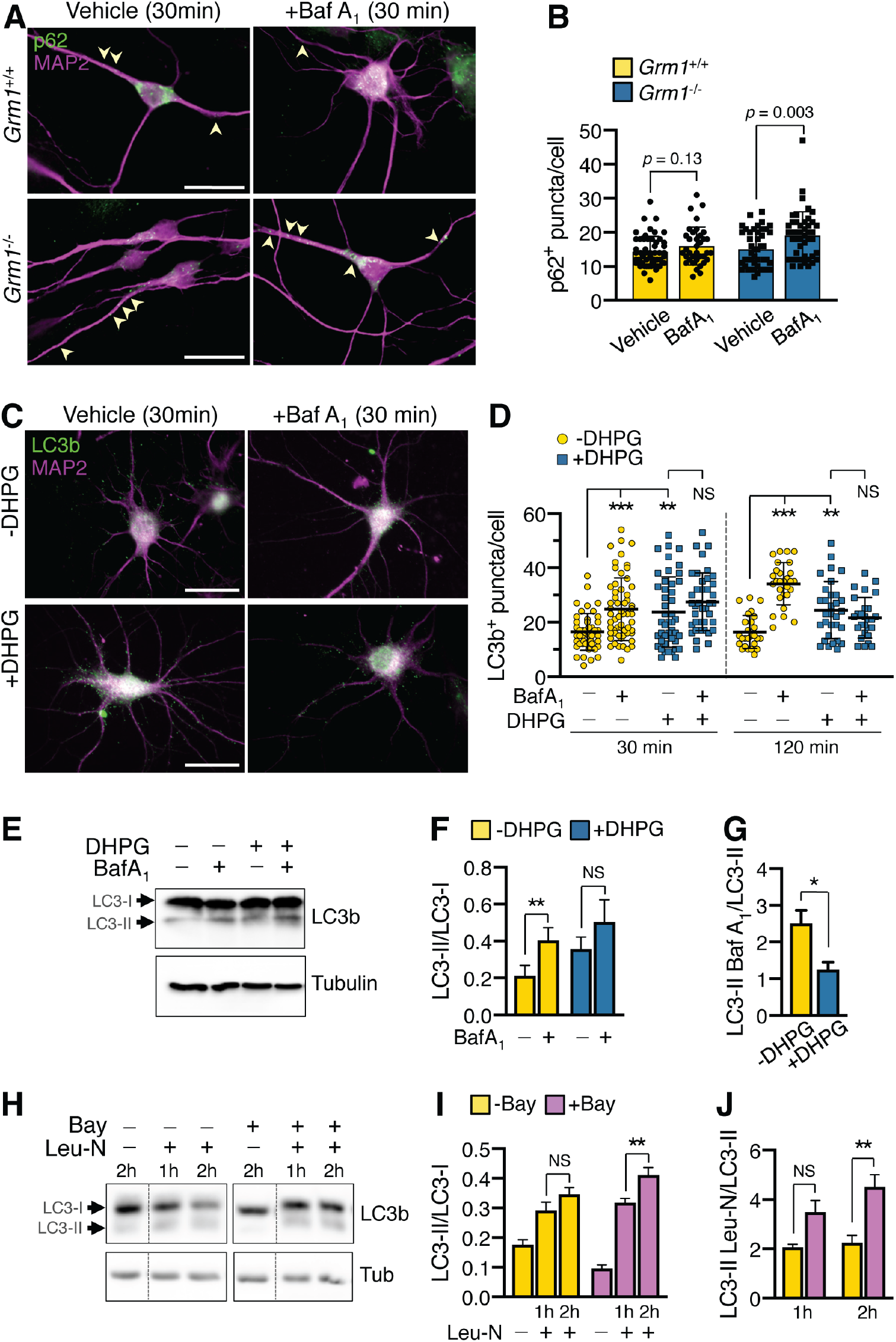
MGlu1 receptor represses autophagy flux in neurons. **(A)** Images of DIV10 mouse hippocampal neurons treated with vehicle or bafA_1_ and labeled with p62 and MAP2; scale bars 25 μm. Arrowheads point to p62^+^ puncta in dendrites (**B**) Quantification of images in (A); each symbol corresponds to p62^+^ puncta in the somatodendritic region of individual neurons. Grm1^+/+^ vehicle n=53, bafA_1_ n=36; Grm1^−/−^ n=45, bafA_1_ n=66 neurons from two independent litters, unpaired t-test. (**C**) Representative images of LC3b and MAP2 in DIV19 rat hippocampal neurons treated with DHPG or vehicle with or without bafA_1_; scale bars 35 μm. (**D**) Quantification of images in (C): each symbol corresponds to LC3b^+^ puncta in the somatodendritic region of each cell (mean±SD). LC3b^+^ at 30 min: −DHPG n=46, +DHPG n=45, −DHPG/bafA_1_ n=58, +DHPG/bafA_1_ n=39. LC3^+^ at 120 min: −DHPG n=26, +DHPG n=31, −DHPG/bafA_1_ n=28, +DHPG/bafA_1_ n=27; *p* values (**) 0.002, (***) <0.001, ANOVA with Tukey post-test. (**E**) LC3b and tubulin immunoblots of rat DIV20 cortical neurons treated as indicated followed by recovery (75 min) with or without bafA_1_. (**F**) Quantification of LC3 turnover as LC3-II/LC3-I ratio (mean±SEM); n=5 biological replicates from two experiments, \*\**p*=0.002, paired t-test. (**G**) Rate of LC3-II accumulation in DHPG-treated cells (normalized to tubulin; mean±SEM): *p*=0.01, unpaired t-test. (**H**) LC3b and tubulin immunoblots of rat cortical neurons treated with Bay36-7620 (Bay) with or without leupeptin and NH_4_Cl (Leu-N). (**I**) Quantification of LC3-II/LC3-I ratio: n=8 biological replicates from two independent experiments, \*\**p*=0.004, ANOVA with Tukey post-test. (**J**) Rate of LC3-II accumulation in Bay-treated cells (normalized to tubulin; mean±SEM): *p*=0.001, ANOVA with Tukey post-test.

Next, we tested whether transient activation of the native receptors could alter basal autophagy flux in neurons. Since glutamate stimulates ionotropic and metabotropic glutamate receptors and can be excitotoxic, we used the synthetic agonist (*S*)-3,5-Dihydroxyphenylglycine (DHPG; (Schoepp et al., 1994)) to activate mGlu1 and mGlu5 receptors and the inverse agonist Bay 367620 to specifically block mGlu1 activity (Carroll et al., 2001). First, we used immunofluorescence in mature rat hippocampal neurons at DIV19 – when mGlu1 expression is at peak – to visualize formation and turnover of LC3b^+^ autophagic vacuoles under basal conditions in soma and MAP2-labeled dendrites. DHPG (50 μM) was applied for 15 min followed by washout and recovery in the absence or presence of bafA_1_. In control neurons, LC3^+^ puncta increased significantly with bafA_1_ treatment *(**Figure 2C-D*** and ***Figure 2 – figure supplement 1***) indicating block of autophagosome–lysosome fusion. Application of DHPG in absence of bafA_1_ increased LC3b^+^ puncta at 30 minutes after agonist washout, an effect that was long-lasting as it persisted for 120 min after stimulation (***Figure 2C-D*** and ***Figure 2 – figure supplement 1***). However, addition of bafA1 to DHPG-treated cells did not significantly increase LC3b^+^ puncta at 30 or 120 minutes after stimulation with agonist *(**Figure 2C-D*** and ***Figure 2 – figure supplement 1***). As an alternative strategy, we used immunoblot to measure LC3-II turnover under basal conditions in DIV20 rat cortical neurons treated with DHPG. Whereas in control cells undigested LC3-II increased substantially upon incubation with bafA_1_, stimulation with DHPG reduced the rate of accumulation of LC3-II *(**Figure 2E-G***). To further confirm the contribution of mGlu1 to the repression of autophagy flux, cortical neurons were treated with the mGlu1-selective inverse agonist Bay 36-7620 (10 μM) to block constitutive activity of endogenous receptors. We found that in cells treated with Bay 36-7620, accumulation of LC3-II in the presence of the lysosomal inhibitors leupeptin and ammonium chloride (Singh et al., 2009) increased over time at a rate greater than untreated neurons *(**Figure 2H-J***). Thus, independent genetic and pharmacological evidence indicate that mGlu1 activity limits the flow of constitutive autophagy in neurons.

### MGlu1 receptor interacts with the adaptor protein FEZ1

Thus far independent measures *in vivo* and *in vitro* converge to implicate mGlu1 receptor activity in balancing basal autophagy in neurons. But what are the molecular effectors that could mediate mGlu1 capacity to regulate autophagy? To address this question we used an unbiased screen by yeast two-hybrid to identify mGlu1 interacting partners. We screened a rat brain library with the carboxyl terminus of rat mGlu1 variant b (mGlu1_b_) (Tanabe et al., 1992), which is expressed in pyramidal neurons of cortex and hippocampus (Berthele et al., 1998; Ferraguti et al., 2008) and retrieved the adaptor protein fasciculation and elongation protein zeta-1 (FEZ1), the mammalian ortholog of UNC-76 in invertebrates (Bloom and Horvitz, 1997; Kuroda et al., 1999) *(**Figure 3 – figure supplement 1***). In vertebrates, FEZ1 is almost exclusively expressed in the brain with prominent expression in cortex and hippocampus (Honda et al., 2004) whereas the closely related FEZ2 is near ubiquitous outside the nervous system (Fujita et al., 2004). We applied orthogonal assays to validate mGlu1 interaction with FEZ1. Immobilized glutathione *S-* transferase (GST) fused to FEZ1, but not GST alone, pulled down MYC-mGlu1_b_ expressed in HEK293 cells validating the interaction *in vitro* (***Figure 3A***). To confirm that the cytoplasmic tail domain of the receptor is sufficient to mediate the interaction with FEZ1, we fused the tail of rat mGlu1_b_ to a reporter derived from a truncated form of vesicular stomatitis virus envelope glycoprotein G (VSV-G) (Francesconi and Duvoisin, 2002). GST-FEZ1 precipitated the VSV-G protein chimera expressed in HEK293 cells (***Figure 3B***) indicating that the receptor tail is sufficient to support interaction with FEZ1. Next, we sought to identify the domains of FEZ1 mediating interaction with the receptor. FEZ1 is a native homodimer (Alborghetti et al., 2010; Lanza et al., 2009): its N-terminus harbors unfolded regions (aa 1-69; 110-229) and participates in dimerization, whereas its carboxyl terminus includes predicted coiled-coil motifs (aa 231-306) that are often associated with scaffolding or vesicle-tethering functions (Truebestein and Leonard, 2016). In pull-down assays, the N-terminal region of FEZ1 spanning aa 1-134 fused to GST was sufficient and necessary for robust precipitation of MYC-mGlu1_b_ expressed in HEK293 cells *(**Figure 3C-D***) whereas precipitation by the carboxyl terminal region spanning aa 223-392 that includes the coiled-coil motifs was marginal *(**Figure 3D***). In co-immunoprecipitation assays, GFP-FEZ1 precipitated MYC-mGlu1b when both co-expressed in heterologous cells (***Figure 3E***) and native FEZ1 co-precipitated with mGlu1b in the hippocampus (***Figure 3F***). Thus three independent measures confirm interaction of the mGlu1 receptor with FEZ1.

**Figure 3.**
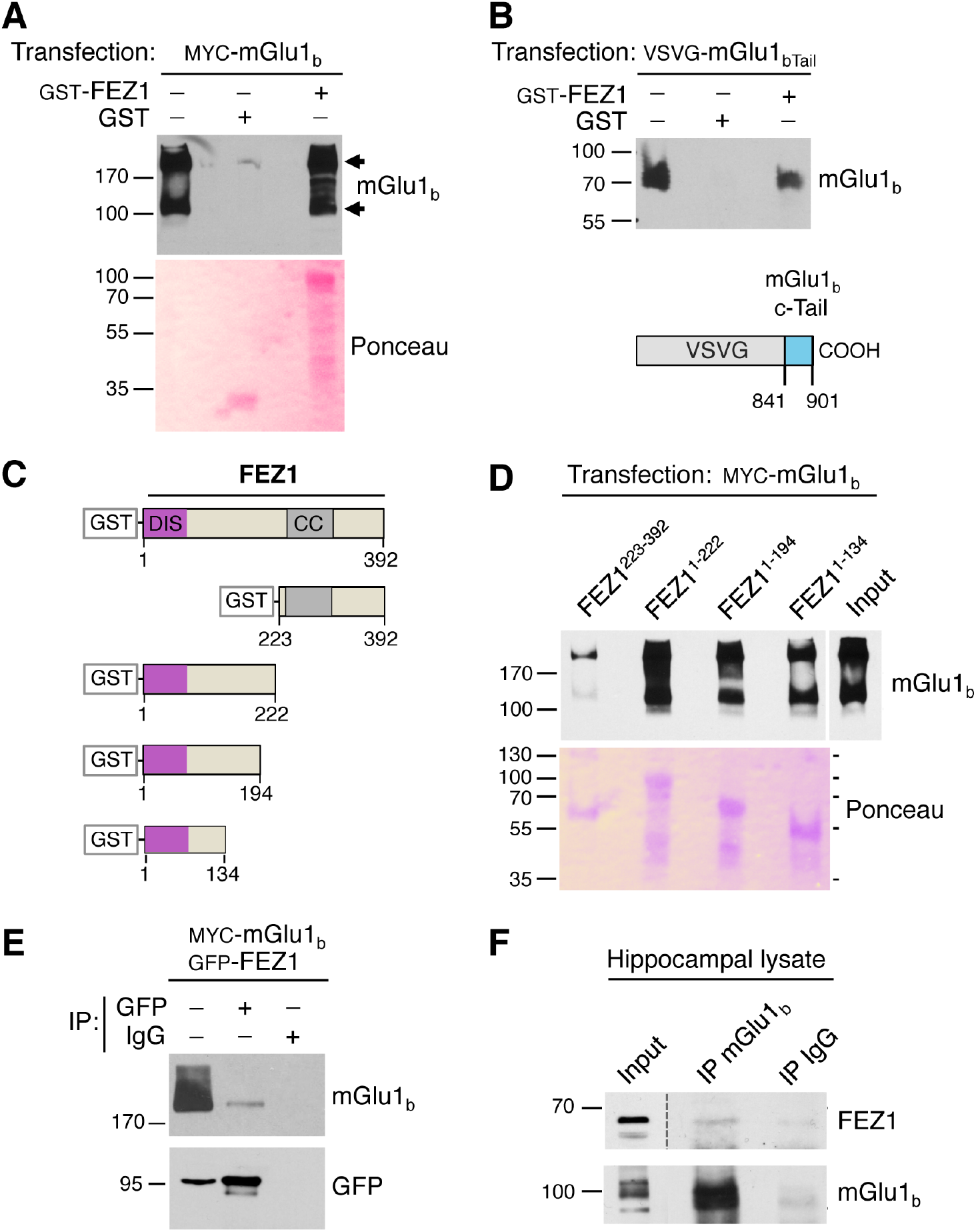
The mGlu1 receptor interacts with the adaptor protein FEZ1. GST-FEZ1 precipitates MYC-mGlu_1b_ from lysates of transfected HEK293 cells. (**A**) Representative immunoblot probed with anti-mGlu_1b_; Ponceau S staining visualizes input proteins in the membrane. Arrowheads point to mGlu1_b_ monomers and dimers; numbers to the left indicate M.W. in kDa. (**B**) The COOH tail of mGlu_1b_ is sufficient for FEZ1 binding. GST-FEZ1 precipitates a protein chimera made of VSV-G fused to the tail of mGlu1b transfected in HEK293 cells: shown is a schematic of the chimeric construct with the mGlu1b tail region indicated in cyan (numbers correspond to residues in rat mGlu1b). (**C-D**) The N-terminus of FEZ1 is sufficient and necessary for interaction with mGlu1_b_. (**C**) Schematic of full-length FEZ1 and deletion fragments to map the regions of FEZ1 that bind mGlu1_b_. (**D**) Immunoblot probed with anti-mGlu1_b_ and membrane stained with Ponceau S: purified FEZ1 fragments fused to GST pull-down mGlu1_b_ expressed in HEK293 cells. (**E**) GFP-FEZ1 co-immunoprecipitates with mGlu1_b_ in transfected cells; mouse IgG served as negative control. GFP-FEZ1 migrates as dimer in SDS-PAGE. (**F**) MGlu1 associates with FEZ1 in the brain. Anti-mGlu1 immunoprecipitates FEZ1 in rat hippocampal lysates; rabbit IgG was used as negative control, immunoblots probed for FEZ1 and mGlu1b.

### FEZ1 supports basal autophagy in neurons

Our findings have uncovered FEZ1 as a novel mGlu1 interacting protein. FEZ1 was previously shown to regulate starvation-induced and basal autophagy in immortalized epithelioid cells (McKnight et al., 2012; Spang et al., 2014). The finding that FEZ1 interacts with mGlu1 suggested that it could play a role in the capacity of the receptor to repress autophagy. However, although FEZ1 is chiefly expressed in the brain, its role in the regulation of autophagy in neurons has not been explored. Thus we first surveyed its role in neuronal autophagy under starvation and basal (nutrient-rich) conditions. For this, we examined the impact of FEZ1 silencing on autophagy flux in neurons incubated in medium without growth-promoting factors, a paradigm that was proposed to enhance autophagy while preserving neural cell health (Young et al., 2009) (but see (Maday and Holzbaur, 2016)). We used pools of four FEZ1-directed small interfering RNAs (siRNAs) or individual siRNAs to induce FEZ1 silencing in rat cortical neurons *(**Figure 4 – figure supplement 1A-B***) in which FEZ1 is expressed at both early and late stages of maturation *in vitro* (***Figure 4 – figure supplement 1A***). In nutrient-depleted medium, DIV12 cortical neurons in which FEZ1 was silenced showed increased LC3-II at steady state compared to control siRNA-treated cells, as determined by immunoblot *(**Figure 4A-B***). Increased LC3-II abundance was not explained by changes in transcription rate given that the relative expression of *Map1lc3b* mRNA measured by RT-PCR was not significantly altered by FEZ1 down-regulation (***Figure 4 – figure supplement 2***). Upon inhibition of lysosomal proteases with leupeptin and ammonium chloride, the rate of accumulation of undigested LC3-II was substantially reduced in cells in which FEZ1 was silenced compared to control siRNA-treated cells *(**Figure 4A,C**)* indicative of decreased autophagy flux. Next, we examined the impact of FEZ1 silencing on basal autophagy in neurons maintained in nutrient-rich medium. We found that FEZ1 downregulation by single siRNAs reduced the rate of accumulation of undigested LC3-II in the presence of lysosomal inhibitors compared to control siRNA-treated cells *(**Figure 4D-F*** and ***Figure 4 – figure supplement 3A-B***). As an independent measure, we used immunolabeling to visualize LC3b in the somatodendritic compartment of rat hippocampal neurons – where FEZ1 is highly expressed (***Figure 4 - figure supplement 4***) – maintained in nutrient-rich medium. Immunolabeled LC3b^+^ puncta were present in both soma and dendrites of control siRNA-treated neurons and were significantly more abundant in FEZ1 siRNA-treated cells *(**Figure 4G-H*** and ***Figure 4 – figure supplement 3C-D***). To survey the digestion of autophagy cargo, we used immunolabeling to visualize the autophagy receptor p62. Labeled p62^+^ puncta were apparent in both control and FEZ1 siRNA-treated cells but more abundant in neurons in which FEZ1 was silenced *(**Figure 4G-H*** and ***Figure 4 – figure supplement 3C-D***). Thus, acute suppression of FEZ1 expression in neurons results in decreased LC3 turnover and accumulation of LC3b^+^ autophagic vacuoles and p62, indicating that FEZ1 supports the progression of autophagy flow in neurons.

**Figure 4.**
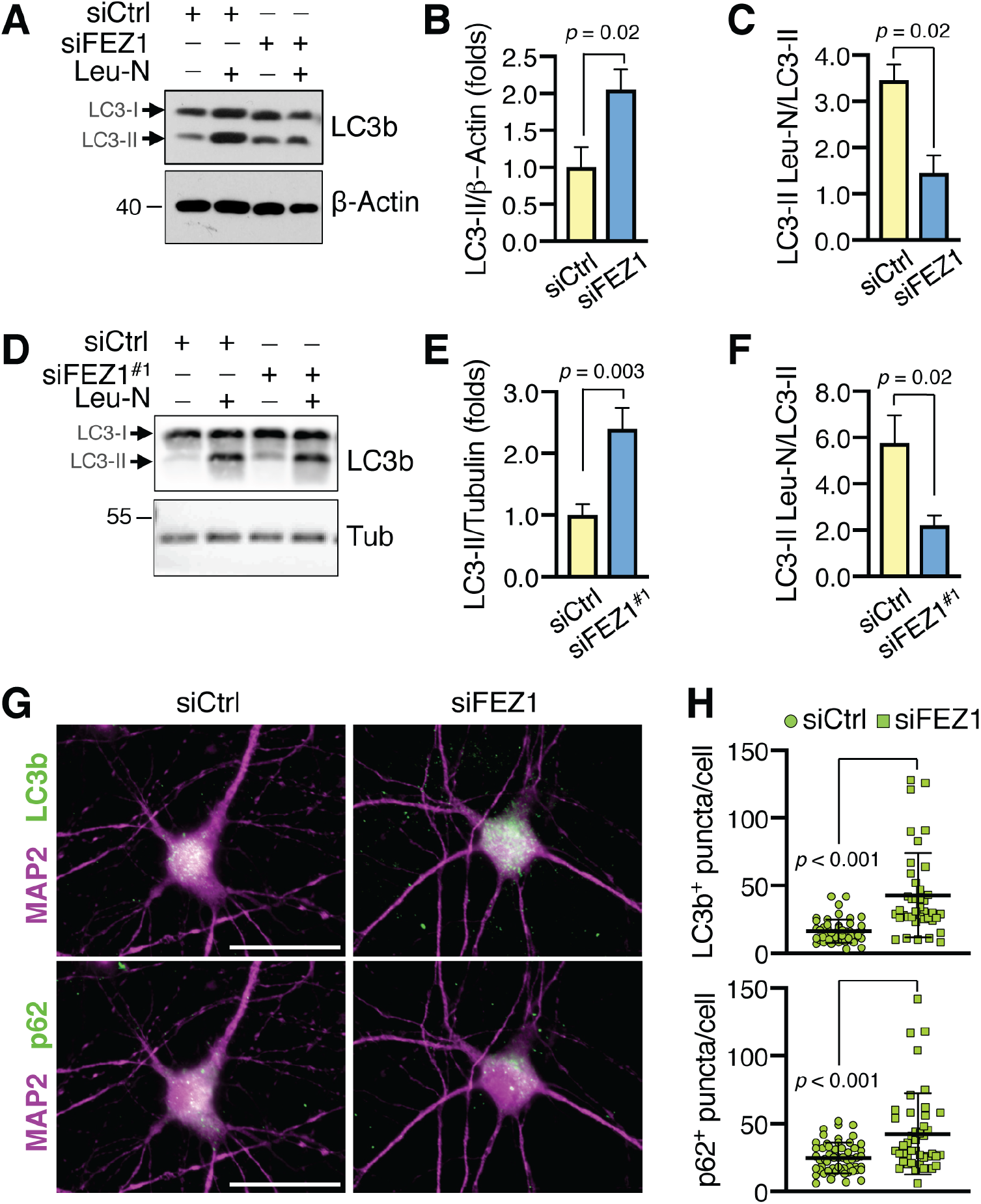
FEZ1 regulates autophagy flux in neurons. (**A**) LC3b and β-actin immunoblots of DIV12 rat cortical neurons transfected with control (siCtrl) or FEZ1 siRNAs (siFEZ1; pool of four siRNAs) and kept in nutrient-depleted medium (90 min) with or without leupeptin and NH_4_Cl (Leu-N). **(B)** Quantification of LC3-II at steady state (normalized to β-Actin; mean±SEM, fold siCtrl): n=6 from three independent cultures, unpaired t-test. (**C**) Rate of LC3-II accumulation (mean±SEM): N=3 experiments, unpaired t-test. (**D**) LC3b and tubulin immunoblots of rat cortical neurons, transfected with control (siCtrl) or individual FEZ1 siRNA (siFEZ1#^1^), kept in nutrient-rich medium with or without leupeptin and NH_4_Cl. (**E**) Quantification of LC3-II at steady state (normalized to tubulin): siCtrl n=7, siFEZ1#^1^ n=6 from two independent cultures, unpaired t-test. (**F**) Rate of LC3-II accumulation (normalized to tubulin): siCtrl n=7, siFEZ1#^1^ n=6, unpaired t-test. (**G**) Images of LC3b (top) and p62 (bottom) in rat DIV12 hippocampal neurons transfected with control or FEZ1 siRNAs (pool of four siRNAs) and kept in nutrient-rich medium; scale bars 35 μm. (**H**) Quantification of images in (G). Each symbol corresponds to LC3b^+^ or p62^+^ puncta in the somatodendrtic compartment of individual neurons. Mean±SD, LC3b^+^ siCtrl n=48, siFEZ1 n=40; p62^+^ siCtrl n=56, siFEZ1 n=44, Mann-Whitney test.

### Interaction with FEZ1 is required for mGlu1-induced repression of autophagy

Although our findings indicate that mGlu1 receptor activity represses constitutive autophagy, the mechanistic link between mGlu1 and the autophagy pathway remained unresolved. The discovery that the mGlu1-binding partner FEZ1 regulates autophagy flow in neurons prompted us to test if interaction with FEZ1 was required for repression of autophagy by the receptor. For this, we took advantage of an anti-FEZ1 antibody that binds the domain that is necessary and sufficient for FEZ1 binding to mGlu1 (aa 1-134), and used it to compete FEZ1 interaction with the receptor in live cells. FEZ1 antibody and control IgG were non-covalently coupled to a cell-penetrating peptide that rapidly shuttles macromolecules into cells (Chariot), and were delivered to rat hippocampal neurons kept in nutrient-rich medium. Autophagy flux was surveyed in the absence or presence of bafA1 by measuring accumulation of the cargo receptor p62 by immunolabeling. As expected, in control neurons incubation with bafA_1_ alone (30 min) resulted in increased p62^+^ puncta in the somatodendritic compartment, and application of DHPG (15 min) prevented accumulation of p62^+^ in bafA_1_-treated cells *(**Figure 5A-B**)*, consistent with inhibition of autophagy flux by receptor activation. Similarly, in neurons that received Chariot-IgG, DHPG prevented bafA_1_-induced accumulation of p62 *(**Figure 5A-B***): in contrast, the effect of DHPG was abolished in neurons transduced with Chariot-FEZ1 antibody, as indicated by increased p62^+^ puncta in the presence of bafA_1_ (**Figure 5*A-B***). Thus occlusion of mGlu1 interaction with FEZ1 abolishes mGlu1 capacity to repress autophagy.

To further validate the requirement for FEZ1 in mGluR-induced repression of autophagy, we used the mRFP/GFP-LC3 reporter to visualize autophagy flux *in situ* (Kimura et al., 2007; Klionsky et al., 2016). In this assay, GFP fluorescence is quenched in the low pH environment of acidified vacuoles whereas mRFP fluorescence is not, thus marking autolysosomes. Rat cortical neurons nucleofected with mRFP/GFP-LC3 were treated with control or FEZ1 siRNAs and switched to nutrient-depleted medium at DIV12 to examine autophagy flux in soma and neurites. After 90 min adaptation, cell were incubated with fresh medium with or without DHPG (50 μM) for 30 min then fixed and imaged. Nucleofected cells displayed numerous mRFP^+^ puncta marking autolysosomes, and sparse mRFP^+^/GFP^+^ puncta consistent with previous reports (Lee et al., 2011)(***Figure 5C-D***). In control siRNA-treated neurons, stimulation with DHPG increased the percentage of mRFP^+^/GFP^+^ puncta, presumably corresponding to non-acidified autophagosomes, compared to vehicle-treated neurons (***Figure 5C-D***). In contrast, in neurons in which FEZ1 was silenced, incubation with DHPG did not significantly alter the relative fraction of mRFP^+^/GFP^+^ puncta compared to control (***Figure 5C-D***). The observed increase in non-acidified autophagosomes is consistent with the capacity of activated receptors to suppress autophagy flux by transient inhibition of autophagosome maturation and/or fusion with acidified vacuoles, a process that is dependent on FEZ1.

**Figure 5.**
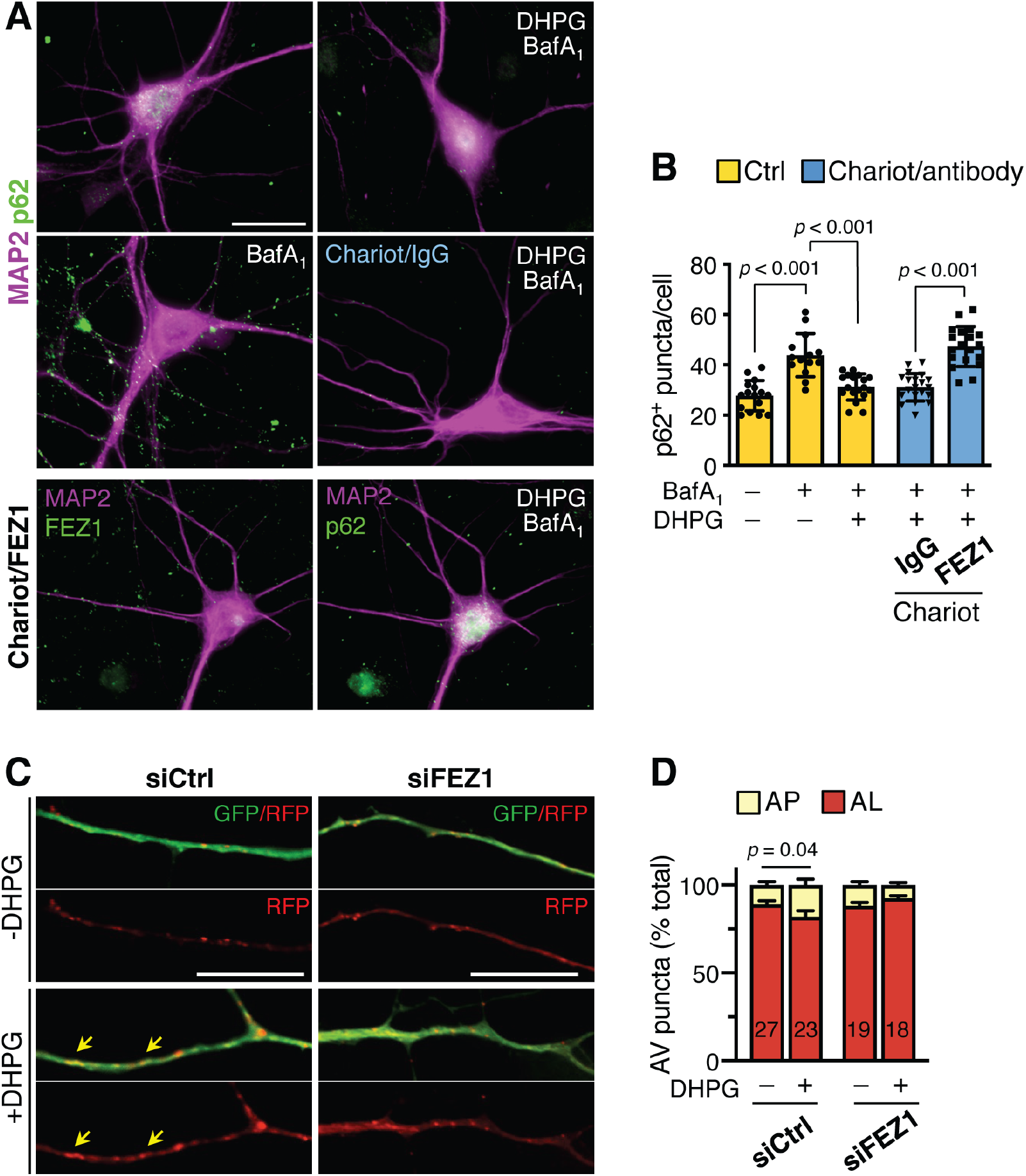
FEZ1 is required for mGluR-dependent repression of autophagy in neurons. (**A**) Images of p62 and MAP2 in rat DIV11 hippocampal neurons untreated or transduced with Chariot peptide coupled to FEZ1 antibody or non-targeting IgG. Neurons kept in nutrient-rich medium were incubated with vehicle or bafA1 with or without DHPG stimulation; scale bar, 25 μm. Uptake of Chariot-FEZ1 antibody (bottom left panel) is confirmed by visualization after addition of corresponding fluorescent secondary antibody only. (**B**) Quantification of images in (A): symbols in the graph correspond to p62^+^ puncta in the somatodendritic compartment of individual neurons. Mean±SD, p62^+^ untreated n=16, bafA1 n=14, DHPG/bafA1 n=18, IgG/DHPG/bafA1 n=20, FEZ1/DHPG/bafA1 n=19, ANOVA with Tukey post-test. (**C**) Images of rat DIV12 cortical neurons expressing mRFP/GFP-LC3 treated with control (siCtrl) or FEZ1 siRNA (siFEZ1; pool of 4 siRNAs) and kept in nutrient-depleted medium. Shown are merged and individual RFP channels; scale bars, 15 μm. (**D**) Quantification of autophagic vacuoles (AV) positive for GFP and RFP (AP, autophagosomes) or RFP only (AL, autolysosomes) as percentage of total AV; numbers in bar graph indicate cells per group from 3 experiments, Mann-Whitney test.

### FEZ1 enables mGlu1 receptor interaction with Ulk1

Our findings indicate that mGlu1 interaction with the adaptor protein FEZ1 is required for its ability to regulate autophagy in neurons. The rapid action of mGlu1 in suppressing autophagy and its dependence on physical interaction with FEZ1, suggested that FEZ1 may form an intermolecular bridge between mGlu1 and the autophagy machinery. Structurally, FEZ1 is a multipronged adaptor protein (Alborghetti et al., 2010) that interacts with multiple partners including the serine-threonine kinase Ulk1, an interaction conserved across phyla (McKnight et al., 2012; Toda et al., 2008). Ulk1 is a critical regulator of autophagy initiation (Mizushima, 2010) (Mercer et al., 2018) but also participates in autophagosome maturation and fusion with lysosomes (Kraft et al., 2012) (Wang et al., 2018), cargo recruitment (Turco et al., 2020) and clearance (Joo et al., 2011). We hypothesized that FEZ1 may form an intermolecular bridge between mGlu1 and Ulk1. To test this, we used immunoprecipitation assays in COS-7 cells and rat brain cortex. In control COS-7 cells, transfected MYC-mGlu1b precipitated FLAG-Ulk1 *(**Figure 6A**)* but silencing of endogenous FEZ1 by stable insertion of a FEZ1 shRNA strongly diminished the co-precipitation of FLAG-Ulk1 with MYC-mGlu1_b_ *(**Figure 6A***). In brain extracts, immunoprecipitation with anti-mGlu1b retrieved endogenous Ulk1 (***Figure 6B***), indicating association between native proteins. Together, these findings provide evidence that mGlu1 can associate with Ulk1 through FEZ1. Ulk1 is regulated by multiple kinases including mTOR, which phosphorylates Ulk1 at Ser^757^ to inhibit its activity (Kim et al., 2011). The finding that mGlu1 can associate with Ulk1, led us to examine if absence of mGlu1 could affect Ulk1 activity by monitoring its phosphorylation by mTOR in the brain of *Grm1*^−/−^ mice. In agreement with previous reports (Tomoda et al., 2004), Ulk1 co-fractionated in soluble and insoluble protein fractions of brain cortices of wild type mice (***Figure 6C***): notably, phosphorylation at Ser^757^ of insoluble Ulk1 and its abundance were reduced in *Grm1*^−/−^ mice compared to wild type *(**Figure 6C-D**)* suggesting alterations in its activity. Based on these observations, we propose that FEZ1 provides a direct physical and functional link between the mGlu1 receptor and the core autophagy machinery.

**Figure 6.**
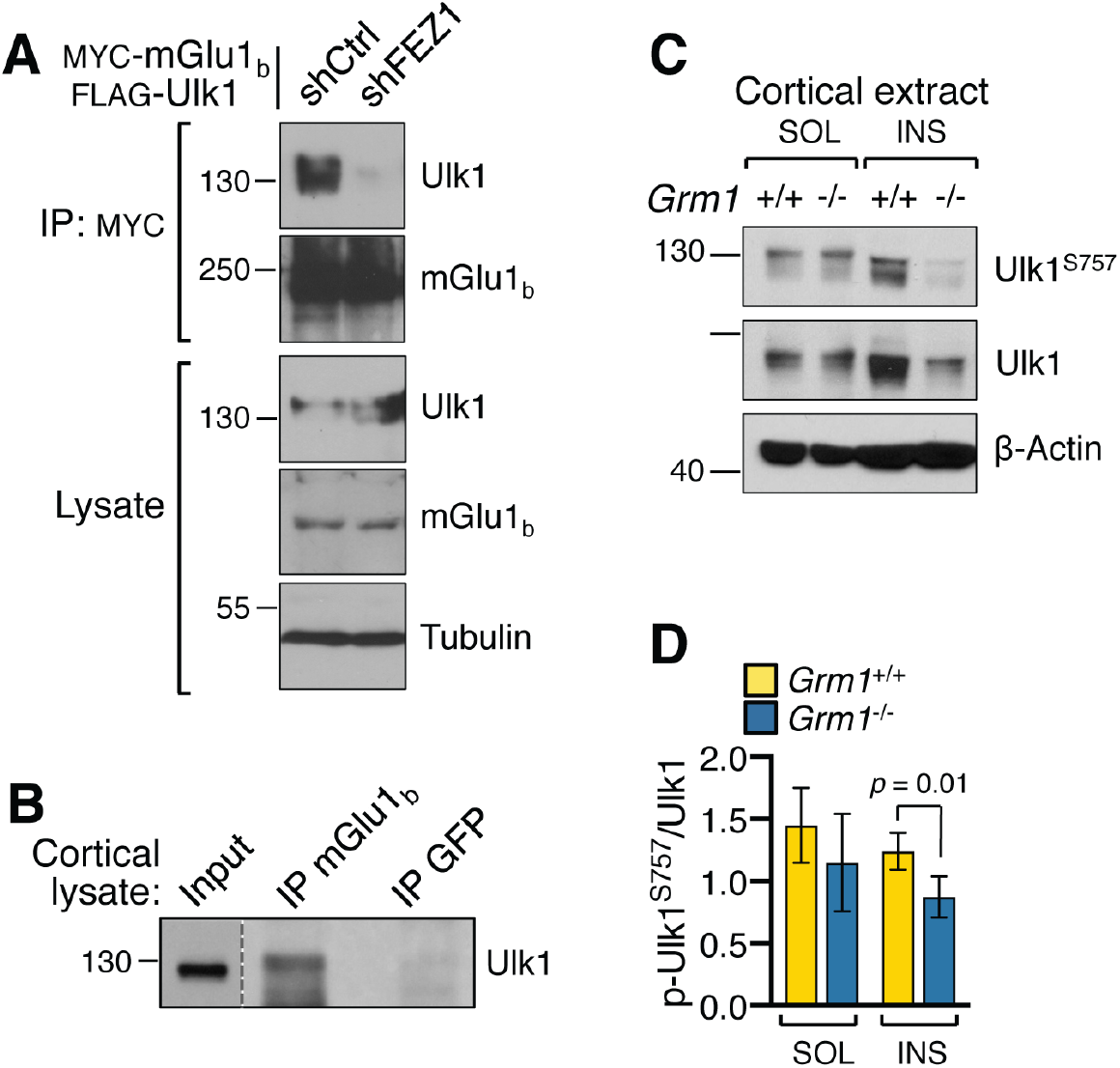
FEZ1 enables MGlu1 association with Ulk1. (**A**) Immunoblot of FLAG-Ulk1 immunoprecipitation by MYC-mGlu1_b_ in COS-7 cells with stably inserted control (shCtrl) or FEZ1 shRNA (shFEZ1) and transfected with the indicated plasmids. (**B**) Immunoblot of Ulk1 immunoprecipitation by anti-mGlu1_b_ in rat brain cortex; anti-GFP was used as negative control. (**C**) Representative immunoblot of total and phosphorylated Ulk1 (Ser^757^) in soluble (SOL) and insoluble (INS) protein fractions of brain cortex from adult wild type and *Grm1*^−/−^ littermates. (**D**) Quantification of Ulk1 phosphorylation at Ser^757^ normalized to total Ulk1. Mean±SEM, SOL *p*=0.53, INS *p*=0.01, N=5 mice per group, paired t-test.

## Discussion

The physiological stimuli that trigger the initiation, fine-tune the progression or curb autophagy in neurons remain little understood. Nutrient deprivation that stimulates autophagy in non-neuronal cells has little impact in the CNS (Maday and Holzbaur, 2016; Mizushima et al., 2004) suggesting that alternative input signals may contribute to the regulation of autophagy in neurons. Here, we provide evidence of a novel role of the mGlu1 receptor in the regulation of autophagy in mammalian neurons and begin to shed insight on underlying mechanisms.

First, we document an imbalance of autophagy in the brain of adult *Grm1*^−/−^ mice. Mutant animals display increased LC3b^+^ puncta and LC3-II but decreased abundance of the autophagy receptors p62 and NBR1, the latter undergoing more efficient lysosomal turnover. Although the data do not identify specific cargo, altered turnover of p62 and NBR1 suggests that selective autophagy of ubiquitinated substrates could be altered. Nevertheless, these observations converge to indicate that global loss of mGlu1 receptor activity in the brain results in hyperactivation of autophagy.

Second, using independent genetic and pharmacological approaches, we show that the mGlu1 receptor represses basal autophagy flux in neurons. Our data indicate that mGlu1 deletion increases the rate of autophagy cargo degradation in the somatodendritic compartment of primary neurons, and that acute inhibition of mGlu1 receptor activity is sufficient to increase the rate of LC3 turnover. Together these observations suggest that mGlu1 receptor activity may inhibit late stages of autophagosome maturation or fusion with degradative vacuoles. However our findings do not rule out potential concomitant effects on autophagosome biogenesis. Autophagosomes are at different stages of maturation in different neuronal compartments and it remains unclear to what extent their biogenesis occurs locally in dendrites and if it is driven by activity (Shehata et al., 2012) (Maday and Holzbaur, 2014). Nevertheless, an emerging scenario is that mGlu1 receptor activity contributes to restrain constitutive neuronal autophagy.

Third, we identify a novel interaction between the carboxyl-terminus of mGlu1 and the N-terminal region of FEZ1, an evolutionarily conserved, neuron-specific adaptor protein.

Fourth, we provide evidence that FEZ1 supports constitutive autophagy in mammalian neurons. FEZ1 and its ortholog UNC-76 interact with kinesin-1 (Blasius et al., 2007; Chua et al., 2012; Gindhart et al., 2003; Toda et al., 2008). In invertebrates, UNC-76 suppression perturbs axonal transport causing formation of aggregates of selected proteins (Chua et al., 2012; Gindhart et al., 2003; Toda et al., 2008) and abnormal clustering of autophagosomes (Chua et al., 2012). A potential scenario is that in mammalian neurons FEZ1 may participate in autophagosome transport and acidification *via* fusion with late endosomes/lysosomes (Lee et al., 2011; Maday et al., 2012). We show that FEZ1 is enriched in the somatodendritic compartment of mature neurons. Although at present direct evidence of progressive autophagosome acidification is limited to axons, recent findings have shown that motile acidified vacuoles are present in dendrites (Goo et al., 2017; Shehata et al., 2012; Winckler et al., 2018) suggesting that an analogous mechanism may operate in dendrites.

Fifth, we show that FEZ1 is required for mGlu1 receptor ability to repress autophagy flux. Competition of FEZ1 interaction with the mGlu1 receptor with blocking antibodies prevents receptor-induced suppression of autophagy cargo degradation, indicating that physical interaction with the FEZ1 adaptor is an essential underpinning of mGluR actions in the regulation of autophagy. FEZ1 forms an antiparallel-oriented dimer (Lanza et al., 2009) (Alborghetti et al., 2013) stabilized by a disulfide bridge (Alborghetti et al., 2010) that imposes a separation between its unfolded N-terminal region and the distal coiled coil domain. This organization endows FEZ1 with properties of a multipronged hub through which mGlu1 could be brought into proximity of other FEZ1 binding partners (Blasius et al., 2007; Chua et al., 2012; Kuroda et al., 1999) that include autophagy adaptors and core components of the autophagy machinery such as NBR1 (Whitehouse et al., 2002) and Ulk1 (McKnight et al., 2012; Toda et al., 2008), respectively. The capacity of FEZ1 to function as protein hub for autophagy is supported by our observation that it enables the association of the mGlu1 receptor with Ulk1. Such association may in part contribute to mGlu1-dependent regulation of constitutive autophagy as suggested by the compromised Ulk1 phosphorylation by mTOR in mGlu1 knockout mice. Reduced phosphorylation of Ulk1 at Ser^757^ and the observed reduced abundance of insoluble Ulk1 could both indicate defects in Ulk1 inactivation (Kim et al., 2018; Liu et al., 2016) (Driessen et al., 2015; Raimondi et al., 2019). However, whether this occurs as a direct consequence of failed mGlu1 signaling to Ulk1, or it arises from potential compensatory homeostatic changes due to upregulated autophagy in *Grm1*^−/−^ mice remains to be determined. Nevertheless, our observations provide evidence that interaction with FEZ1 can link the mGlu1 receptor to the core autophagy machinery.

Autophagy is a highly regulated, dynamic pathway and new functions of established core machinery components, *e.g.* Ulk1, as well as new players are being uncovered. Given the hub properties of FEZ1, it is conceivable that during the course of autophagy flow FEZ1 may provide a transient physical and functional link for mGlu1 to additional components of the autophagy machinery.

Autophagy in vertebrates contributes to the establishment of neuronal connectivity (Dragich et al., 2016), presynaptic function (Hernandez et al., 2012; Vijayan and Verstreken, 2017), pruning of dendritic spines (Lieberman et al., 2020; Tang et al., 2014), synaptic plasticity (Nikoletopoulou et al., 2017) and memory formation (Glatigny et al., 2019; Hylin et al., 2018; Pandey et al., 2020). Moreover, emerging evidence indicates neuronal type–specific reliance on autophagy for dendritic morphogenesis and control of excitability (Lieberman et al., 2020). At present, we know little on the contribution of neurotransmitter receptors to the regulation of neuronal autophagy. GPCRs were recently shown to regulate autophagy in non-neuronal cells (Zhang et al., 2015) and in tissue outside the CNS (Wauson et al., 2014; Wauson et al., 2012). Here we show that in the CNS the mGlu1 receptor plays a critical role in balancing autophagic activity, a function likely conserved through evolution (Kang and Avery, 2009) and potentially shared by other mGluRs since mGlu5 receptor inhibition reduces the burden of huntingtin aggregates in a mouse model of Huntington’s disease *via* autophagy (Abd-Elrahman et al., 2017). Deregulated mGlu1 receptor signaling is implicated in neurodevelopmental disorders including schizophrenia and autism spectrum disorders (Foster and Conn, 2017). Future studies will need to address to what extent mGlu1-induced repression of autophagy contributes to receptor functions in health and disease.

## Methods

### Animals

The *Grm1* mutant mouse strain used is described in (Conquet et al., 1994). Experimental animals were fed *ad libitum* and mutant mice co-housed with wild type littermates. Animals of both sexes were used for experiments. Newborn *Sprague Dawley* rat pups of both sexes were used for primary cultures. All procedures involving animals were carried out according to protocols approved by the Albert Einstein College of Medicine Institutional Animal Care and Use Committee and in accordance with the Guide for the Care and Use of Laboratory Animals by the United States PHS.

### Cell lines and neuronal cultures

HEK293 and COS-7 cells (ATCC) were maintained in DMEM supplemented with 10% fetal bovine serum, 1% non-essential amino acids, 100 U/ml penicillin and 100 μg/ml streptomycin. Clonal cells were selected with puromycin (5 μM). Cortical and hippocampal neurons from newborn rat (P1) pups were plated onto poly-L-lysine coated coverslips or multi-well culture plates. Rat neurons were maintained in serum-free medium of Neurobasal A, 2% B27 supplement, 2 mM GlutaMax (all from Gibco), 37 mM Uridine and 27 mM 5-Fluoro-2-deoxyuridine (Sigma Aldrich). Mouse hippocampal neurons were harvested from P4 pups, that were genotyped by PCR before plating, and maintained in serum-free medium of Neurobasal A, 2% NeuroCult SM1 supplement (STEMCELL Technologies), 2 mM GlutaMax.

### Transfection and RNAi

HEK293 and COS-7 cells were transfected with Lipofectamine 2000 (Invitrogen) according to manufacturer’s specifications. Neurons were transfected before plating by nucleofection (Lonza) as described (Francesconi et al., 2009). RNAi was carried out as previously described (Kalinowska et al., 2015) using Accell siRNAs (Dharmacon). The siRNAs were applied at 1 μM final concentration to DIV 7/8 neurons in culture medium and maintained for 4 days. The following siRNAs were used: rat FEZ1 SMARTpools, rat FEZ1 siRNA A-094375-16, rat FEZ1 siRNA A-094375-14, non targeting control D-001910.

### Antibody delivery with Chariot

Chariot peptide (Active Motif, Carlsbad CA) was coupled to rabbit IgG and rabbit anti-FEZ1 antibody (ThermoFisher PA590412) following manufacturer’s recommendations. Briefly, to couple antibodies to the Chariot peptide, 2 μl of Chariot were combined with 1 μg of antibody in PBS, gently mixed and incubated for 30 min at room temperature. After collecting the culture medium, neurons were rinsed once with PBS and fed fresh medium to which Chariot/antibody complex was added. Cells were incubated (37°C, 5% CO2) with gentle rocking for 1 hr after which the previously collected culture medium was added back to the cells for 4 hr.

### RT-PCR

First-strand cDNA was prepared directly from cells and tissue using SuperScript III CellsDirect cDNA Synthesis System (Invitrogen). Briefly, neurons in 24-well plates were incubated with trypsin for 5 min and suspended in medium of DMEM, 10% FBS, 2 mM GlutaMAX. Cells were gently washed with PBS, suspended in PBS and counted: 2 μl of the adjusted cell suspension was transferred to 10 μl of lysis buffer containing RNAseOUT. Cell lysis, DNA digestion, and 1^st^ strand synthesis with oligodT were according to manufacturer’s instructions. For tissue, small explants of brain cortex in RNALater (Life Technologies) from 8-12-month old mice were disrupted with a needle and spun at 1,800 rpm at 4°C for 5 min. The cell pellet was suspended in PBS and 2 μl used for 1^st^ cDNA synthesis. Synthesized products were quantified with nanodrop and equal amounts used in the PCR reactions. Primers used were: mouse LC3b-Fw 5’-AAGAGTGGAAGATGTCCGGC-3’, LC3b-Rv 5’-ACTTCGGAGATGGGAGTGGA-3’, p62-Fw 5’-AGATGCCAGAATCGGAAGGG-3’, p62-Rv 5’-GAGAGGGACTCAATCAGCCG-3’, mouse/rat GAPDH-Fw 5’-ACCACAGTCCATGCCATCAC-3’, GAPDH-Rev 5’-TCCACCACCCTGTTGCTGTA-3’, Rat LC3b-Fw 5’-CGGAGCTTCGAACAAAGAGTG-3’, LC3b-Rv 5’-ACCATGCTGTGCCCATTCAC-3’.

### Yeast two-hybrid

The screen of a rat brain cDNA library was carried out with the GAL4-based MATCHMAKER system (Clontech) and all procedures followed the system protocols. The cDNA library made in the pACT2 vector carrying the GAL4 DNA-AD was transformed in Y187 *(MATα)* yeast cells (1.5×10^6^ independent clones). pACT2 encodes the *LEU2* gene that allows growth in synthetic media lacking leucine (Leu^−^). The bait (BD:mGlu1_Tail_) was generated in the pAS2-1 vector by joining GAL4 DNA-BD in frame to the mGlu1_b_ C-terminus with sequence: KPERNVRSAFTTSDVVRMHVGDGKLPCRSNTFLNIFRRKKPGAGNAKKRQPEFSPSSQCPSAHAQL. pAS2-1 encodes the *TRP1* gene that allows growth in media lacking tryptophan (Trp^−^). The bait was transformed in AH109 yeast cells (*MATa*) and did not induce auto-activation of reporter genes *lacZ* and *HIS3.* The library was screened by interaction-mating *(MATa vs. MATa):* mated cells were plated on medium lacking leucine, tryptophan, histidine and adenine. Of three millions clones screened, 126 diploids grew under selection and reduced to 34 after re-screening for false positives. Plasmid DNA was isolated and analyzed by restriction digest with EcoRI and XhoI to determine insert size. To confirm selective interaction with the bait, individual plasmids were re-transformed in Y187 cells and mated with AH109 or Y190 (*MATa*) cells expressing either the BD:mGlu1_Tail_ bait or the BD:laminC fusion (pLAM5) to eliminate clones interacting with unrelated proteins.

### GST fusion proteins and pull-down assays

Purification of GST fusion proteins expressed in *E. Coli* BL21(D3) and pull-down assays with cell lysates were carried out as described (Kalinowska et al., 2015). Each pull-down used 100 pmol of purified recombinant protein immobilized onto glutathione-agarose beads and 2 mg of cell lysate.

### Immunoprecipitation and Western blot

For immunoprecipitation, adherent cells were harvested on ice in lysis buffer (20 mM Tris-HCl, pH 7.5, 150 mM NaCl, 5 mM EDTA, 1% Triton X-100, 0.5% sodium deoxycholate), briefly sonicated and incubated on ice 15 min. After centrifugation at 14,000 rpm for 15 min the supernatant was recovered and pre-cleared for 10 min with Protein G-coupled magnetic beads (Dynabeads, Life Technologies). The pre-cleared lysate was incubated for 16 hr at 4°C with antibody coupled to the magnetic beads; the immunocomplex was washed once with lysis buffer, 3 times with wash buffer (20 mM Tris-HCl, pH 7.5, 150 mM NaCl, 5 mM EDTA, 0.5% Triton X-100) and once in wash buffer without detergents. The beads were rinsed once in PBS and the immunocomplex eluted in sample buffer. Immunoprecipitation from brain tissue lysates was performed as previously described (Kalinowska et al., 2015). Briefly, dissected cerebrum or hippocampus (P10) from rat was homogenized on ice in a buffer of 10 mM Tris-HCl, 5 mM EDTA, 320 mM sucrose (pH 7.4) with cocktails of protease and phosphatase inhibitors. The homogenate was spun at 800 x *g* for 10 min and resulting supernatant at 10,000 x *g* for 15 min. For immunoprecipitation, pellet and supernatant were equilibrated to 50 mM Tris-HCl (pH 7.4), 150 mM NaCl, 1 mM EDTA with 1% Triton X-100 and 0.5% sodium deoxycholate. Western blot analysis and detection with horseradish peroxidase-conjugated secondary antibodies and ECL was carried out according to standard protocols.

### Tissue fractionation

Freshly microdissected brain cortices were homogenized with a Dounce homogenizer in ice-cold lysis buffer (50 mM Tris-Cl pH 7.42, 150 mM NaCl, 1% NP-40, 1% (w/v) sodium deoxycholate, 0.1% SDS) with cocktails of protease and phosphatase inhibitors and incubated with agitation for 60 min at 4°C. The homogenate was centrifuged at 21,000 *g* for 30 min at 4°C and the supernatant collected (soluble fraction). The pellet was washed with lysis buffer, centrifuged for 5 min, dissolved in 2% SDS in lysis buffer, sonicated, incubated with agitation for 60 min and centrifuged at 21,000 *g* for 10 min (insoluble fraction).

### Flux assays

Flux assays in cortical or hippocampal neurons were carried out in nutrient depleted (DMEM) or nutrient-rich medium (Neurobasal A, 2 mM GlutaMAX, 2% B27 supplement). Two different sets of lysosomal inhibitors were used: 200 μM leupeptin together with 20 mM NH_4_Cl or bafilomycin A1 (100 nM). After rinsing with corresponding fresh medium cells were incubated at 37°C, 5% CO_2_ for indicated times with medium with vehicle or inhibitors. For agonist stimulation, after rinsing with medium cells were incubated with vehicle or with 50 μM S-DHPG (Tocris) for 15 min at 37°C, rinsed with medium and then incubated with fresh medium with or without inhibitors for indicated times. For treatment with antagonist, cells were rinsed and incubated for indicated times with fresh medium with either vehicle or 10 μM BAY 36-7620 (Tocris) in the presence or absence of lysosomal inhibitors. At the end of incubation time, cells were placed on ice, rinsed with ice-cold PBS, scraped off in cold RIPA buffer with cocktails of protease and phosphatase inhibitors and processed for protein analysis. For *ex vivo* flux assays (Esteban-Martinez and Boya, 2015), mouse brain cortices were harvested in ice-cold dissection medium immediately after euthanasia. Finely chopped tissue explants were equally divided into pre-warmed medium (75% MEM, 25% HBSS, 2 mM glutamine) with or without 200 μM leupeptin and freshly made 20 mM NH_4_Cl and incubated at 37°C, 5%CO_2_ for 60 min with occasional swirling. Tissue was transferred to microcentrifuge tubes, washed twice with ice-cold PBS, briefly spun to collect it at the bottom of the tube and supernatant discarded. Tissue was suspended in lysis buffer with a cocktail of protease and phosphatase inhibitors and disrupted on ice with a Dounce homogenizer. Homogenate was incubated with rotation at 4°C for 15 min and processed for immunoblot analysis.

### Immunofluorescence

Cells were washed two times in PBS for 5 min and fixed with 4% paraformaldehyde for 10 min. Cells were permeabilized with 0.1% Triton X-100 in PBS for 15 min and blocked for 60 min at room temperature with 5% BSA or 5% normal serum; primary antibodies diluted in blocking solution were incubated overnight at 4°C. After three washes with PBS, cells were incubated for 60 to 90 min at room temperature with fluorophore-conjugated secondary antibodies, washed three times with PBS and mounted with Prolong (Invitrogen). For brain tissue labeling, coronal sections (25-30 μm thick) were cut with a vibratome from tissue post-fixed in 4% paraformaldehyde overnight. Sections were washed three times (10 min) in PBS, once in PBS with 0.01% Triton X-100 (15 min) and blocked with 10% BSA for 30 min at room temperature. The following primary antibodies were applied for 24 h at 4°C: anti-LC3b (1:100; Cell Signaling Technology), anti-p62 (1:400; BD Transduction Laboratories), anti-MAP2 (1:500; Phosphosolution). Sections were washed and incubated with secondary antibodies as above, washed three times for 5 min and mounted with Prolong antifade with DAPI. Epifluorescence was imaged with 40× (NA = 1.3) or 60× (NA = 1.35) oil objectives mounted on an Olympus IX81 microscope equipped with digital CCD ORCA-R2 camera (Hamamatsu). Confocal images were acquired with Olympus Fluoview 500 Confocal Scanning Microscope. Image analysis was performed with Fiji-Image J (NIH). After background subtraction, neuronal soma were identified by MAP2 overlay: puncta were counted with the Cell_Counter plugin and validated with a secondary analysis using threshold adjustment and the Analyze_Particle plugin for automated counts.

### Antibodies

Goat polyclonal anti-GAPDH (GenScript), chicken polyclonal anti-MAP2 (EnCor Biotech; Phosphosolutions); mouse monoclonal antibodies anti-p62 (BD Transduction Laboratories); anti γ-tubulin (Sigma Aldrich), anti-β-actin (Sigma Aldrich), anti-FEZ1 (Sigma Aldrich), anti-GM130 (BD Bioscience) anti-Beclin1 (BD Bioscience), anti-myc tag (Cell Signaling Technology); rabbit polyclonal antibodies anti-LC3b (Novus Biologicals), anti-mGlu1_b_ (in house; (Mende et al., 2016)), anti-FEZ1 (Sigma Aldrich; ThermoFisher), anti-GFP (Santa Cruz Biotech); from Cell Signaling Technology anti-FEZ1, anti-Ulk1, anti-phospho-Ulk1^S757^, anti-NBR1, anti-LC3b.

### Plasmids

We generated the following plasmids by standard cloning techniques: rat GFP-FEZ1, GST-FEZ1, GST-FEZ1(1-134), GST-FEZ1(1-194), GST-FEZ1(1-222), GST-FEZ1(223-392), BD:mGlu1_Tail_ (pAS2-1 vector, see yeast 2-hybrid). The following plasmids were also used: rat MYC-mGlu1b (with 3 tandem myc tags) and VSVG-mGlu1bTail (Francesconi and Duvoisin, 2002), FLAG-Ulk1 (Addgene; Reuben Shaw lab), pGEX-4T-2 (GE Healthcare), pGIPZ control shRNA, and pGIPZ FEZ1 shRNA (Einstein core facility), mRFP/GFP-LC3 (gift from JD MacMicking, Yale University).

### Statistics

Statistical significance was determined by Student’s *t*-test or Mann-Whitney test in pairwise comparison and ANOVA for multiple groups with *p* values <0.05 considered significant.

## Acknowledgments

Supported by NIMH MH108614 and an early award from NARSAD (now Brain & Behavior Research Foundation). We acknowledge the assistance of the Neural Cell Engineering and Imaging Core of the Einstein Rose F. Kennedy Intellectual and Developmental Disabilities Research Center supported by NICHD U54 HD090260. We thank Ana Maria Cuervo for advice and reagents.

---

**Figure 1 – figure supplement 1.**
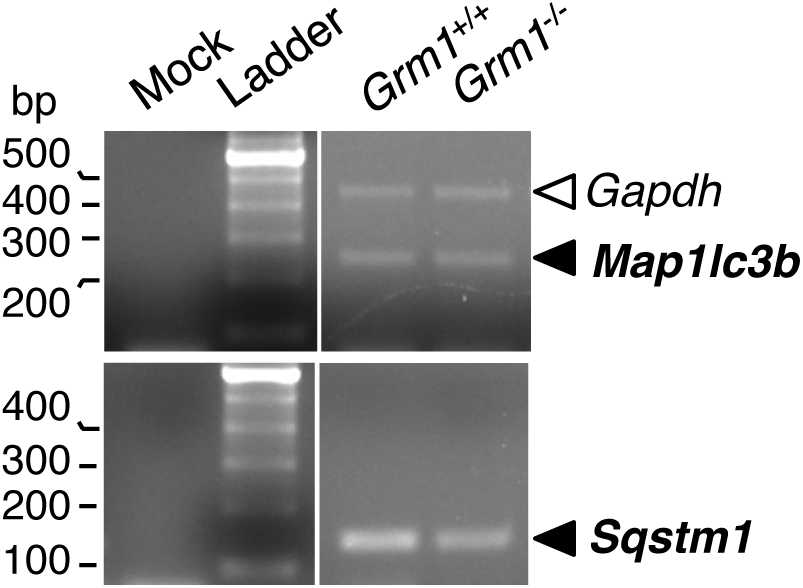
Expression of *Map1lc3b* (LC3b) and *Sqstm1* (p62) mRNA is not altered in the brain cortex of adult *Grm1*^−/−^ mice. Agarose gels with RT-PCR amplicons from total RNA. Relative band intensities (mean±SEM) of *Map1lc3b* and *Sqstm1* were normalized to *Gapdh. Map1lc3b: Grm1^+/+^* 0.54±0.21, *Grm1*^−/−^ 0.50±0.29, *p*=0.7782, N=4 littermates per group, two-tailed paired t-test. *Sqstm1:Grm1^+/+^* 0.76±0.56, *Grm1*^−/−^ 0.40±0.27, *p*=0.3204, N=4. Mock, no 1^st^ strand.

**Figure 2 – figure supplement 1.**
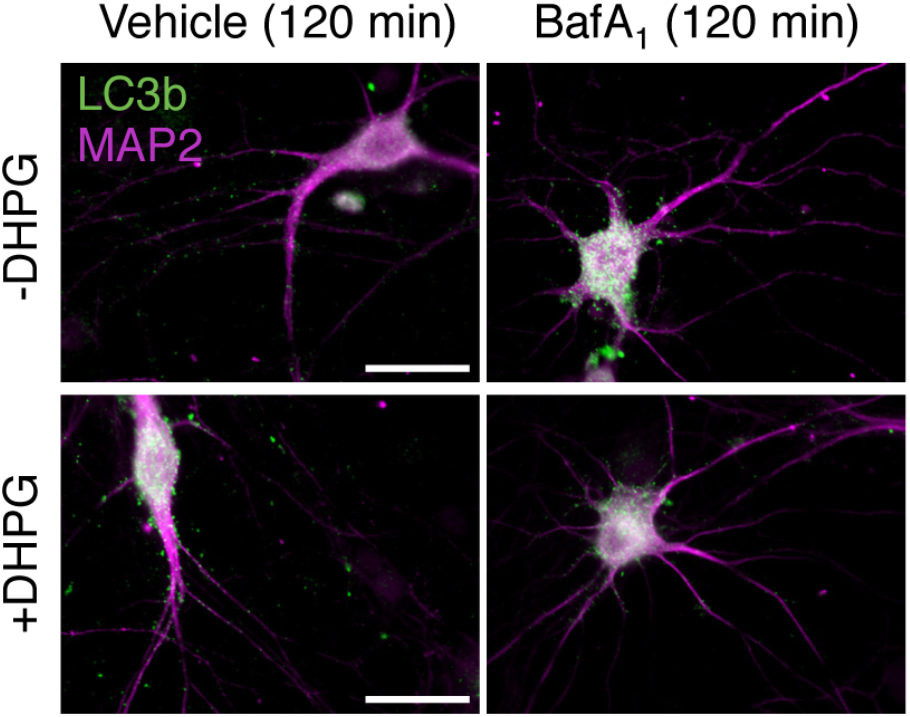
Stimulation with DHPG represses autophagy flow in hippocampal neurons. Representative images of LC3b and MAP2-labeled DIV19 rat hippocampal neurons treated with DHPG or vehicle in absence or presence of bafA_1_ (120 min); scale bars 35 μm.

**Figure 3 – figure supplement 1.**
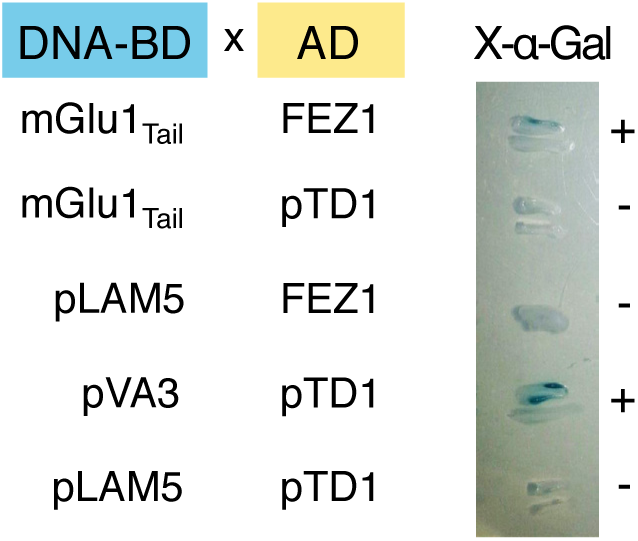
Yeast two-hybrid screen identifies FEZ1 as mGlu1_b_ interactor. A rat brain cDNA library was screened with the bait DNA-BD:mGlu1_Tail_ cloned in pAS2-1 vector containing the mGlu1_b_ COOH-terminus fused in frame with the GAL4 DNA Binding Domain (DNA-BD). Representative X-gal assay illustrates interaction between the DNA-BD:mGlu1_Tail_ bait and FEZ1 clone. MATa and MATα yeast cells were re-transformed with individual plasmids, mated and plated in Leu^−^/Trp^−^ medium with X-gal to visualize activation of reporter *LacZ* gene. Negative controls: DNA-BD:mGlu1_Tail_ (GAL4 DNA-BD, *TRP1*) mated to pTD1 (GAL4 AD:SV40 T antigen, *LEU2);* FEZ1 (GAL4 AD, *LEU2)* mated to pLAM5 (GAL4 DNA-BD:laminC, *TRP1*); pLAM5 mated to pTD1. Positive control: pVA3 (GAL4 DNA-BD:p53, *TRP1*) mated to pTD1.

**Figure 4 – figure supplement 1.**
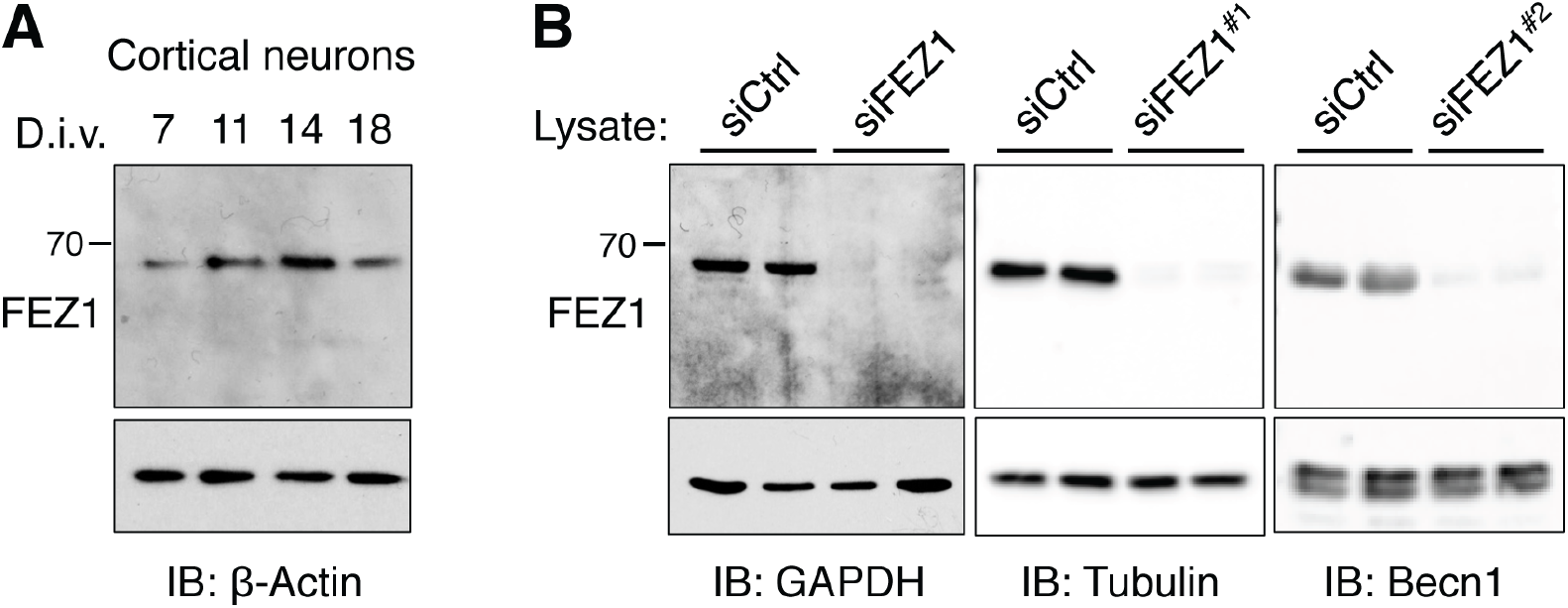
FEZ1 is expressed in primary cortical neurons at different stages of maturation *in vitro.* (**A**) FEZ1 and β-actin immnunoblots of lysates of rat cortical neurons at indicated days *in vitro* (d.i.v). (**B**) Immunoblots of FEZ1 in DIV12 rat cortical neurons transfected with control siRNA (siCtrl) or a pool of four FEZ1 siRNAs (siFEZ1; ~ 95% knockdown) or individual FEZ1 siRNAs (siFEZ1#^1^ and siFEZ1#^2^, ~ 95% and ~ 90% knockdown respectively): loading controls are shown in panels underneath.

**Figure 4 – figure supplement 2.**
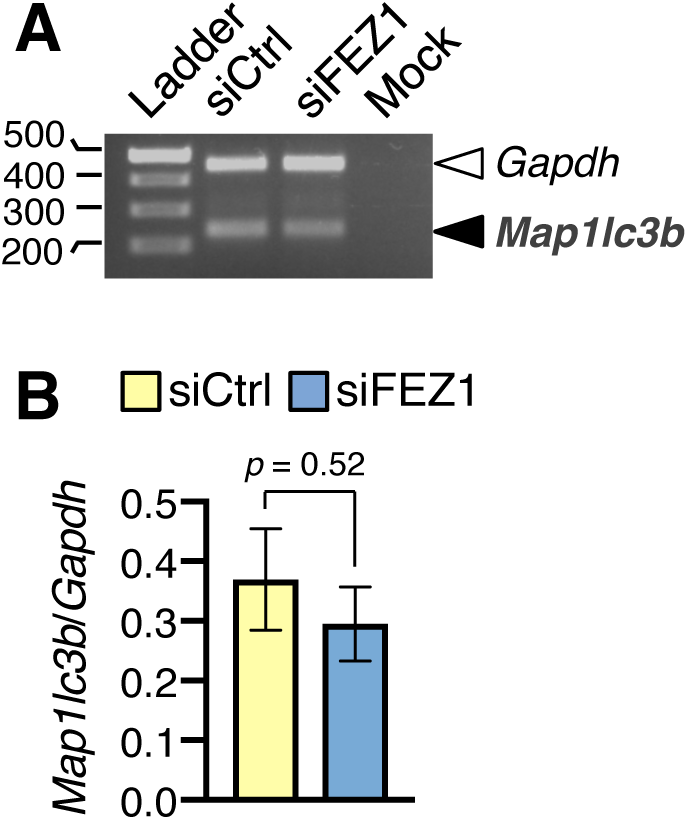
Increased LC3-II at steady state by FEZ1 silencing is transcription-independent. (**A**) *Map1lc3b* (LC3b) mRNA in rat cortical neurons treated with control (siCtrl) or FEZ1 siRNA (siFEZ1; pool of four siRNAs). Agarose gel with *Map1lc3b* and *Gapdh* amplicons from RT-PCR of total RNA; mock, no 1^st^ strand. (**B**) Quantification of *Map1lc3b* expression normalized to *Gapdh.* Mean±SEM, n=3 experiments, unpaired t-test.

**Figure 4 – figure supplement 3.**
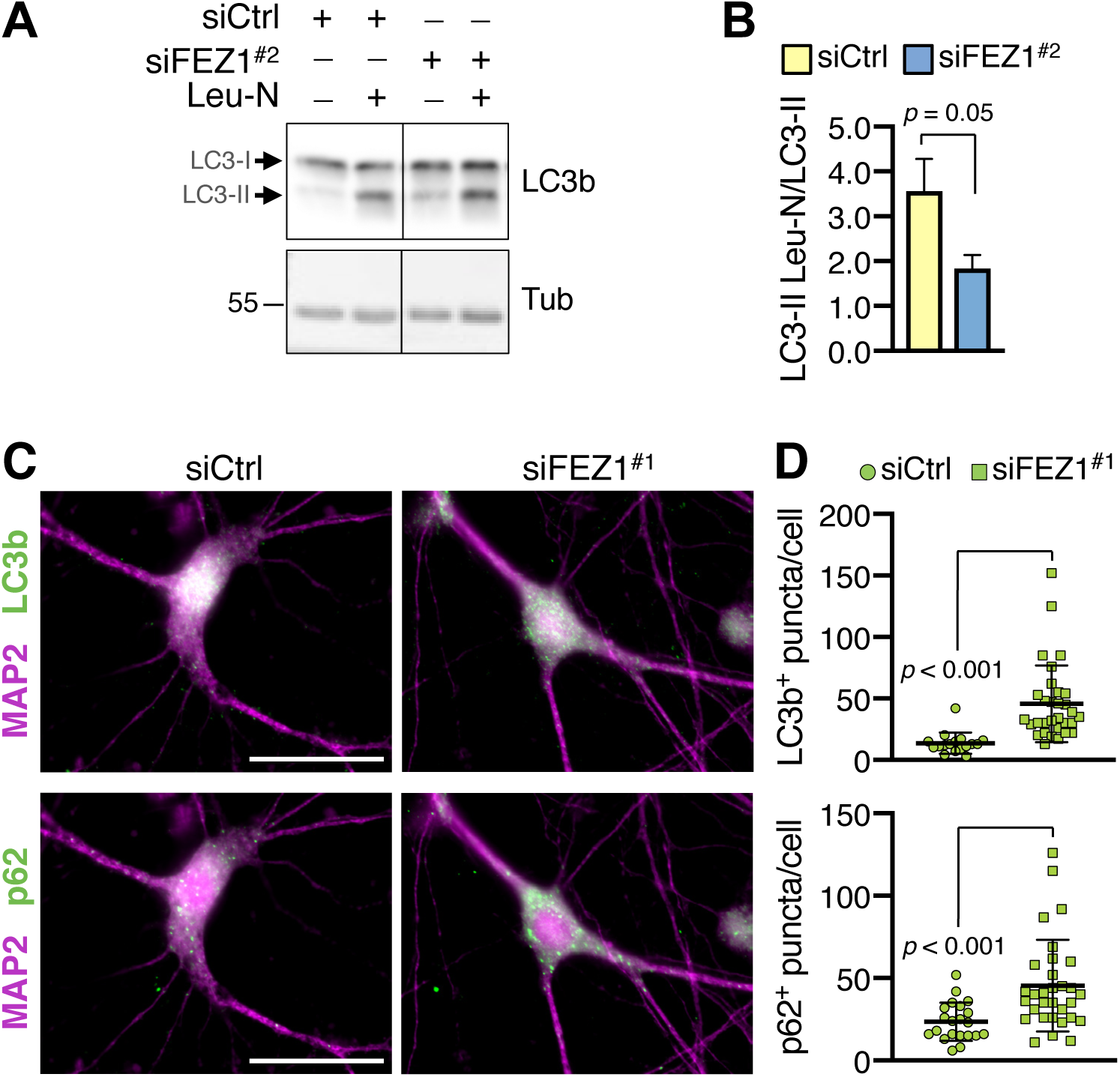
FEZ1 silencing impairs autophagy flux in neurons. (**A**) LC3b and tubulin immunoblots of lysates of rat cortical neurons transfected with control (siCtrl) or individual FEZ1 siRNA (siFEZ1^#2^), kept in nutrient-rich medium with or without leupeptin and NH4Cl (Leu-N). (**B**) Rate of LC3-II accumulation (normalized to tubulin, mean±SEM): n=6 individual knockdown from two independent cultures, unpaired t-test. (**C-D**) FEZ1 silencing increases the density of LC3b^+^ and p62^+^ puncta in rat hippocampal neurons in nutrient-rich medium. (**C**) Images of LC3b (top) and p62 (bottom) in soma and MAP2-labeled dendrites of rat DIV12 hippocampal neurons transfected with control (siCtrl) or individual FEZ1 siRNA (siFEZ1#^1^); scale bars 35 μm. (**D**) Quantification of images in (C); each symbol corresponds to somatodendritic LC3b^+^ or p62^+^ puncta per neuron. Mean±SD, LC3b^+^ siCtrl n=17, siFEZ1#^1^ n=31; p62^+^ siCtrl n=21, siFEZ1#^1^ n=31, Mann-Whitney test.

**Figure 4 – figure supplement 4.**
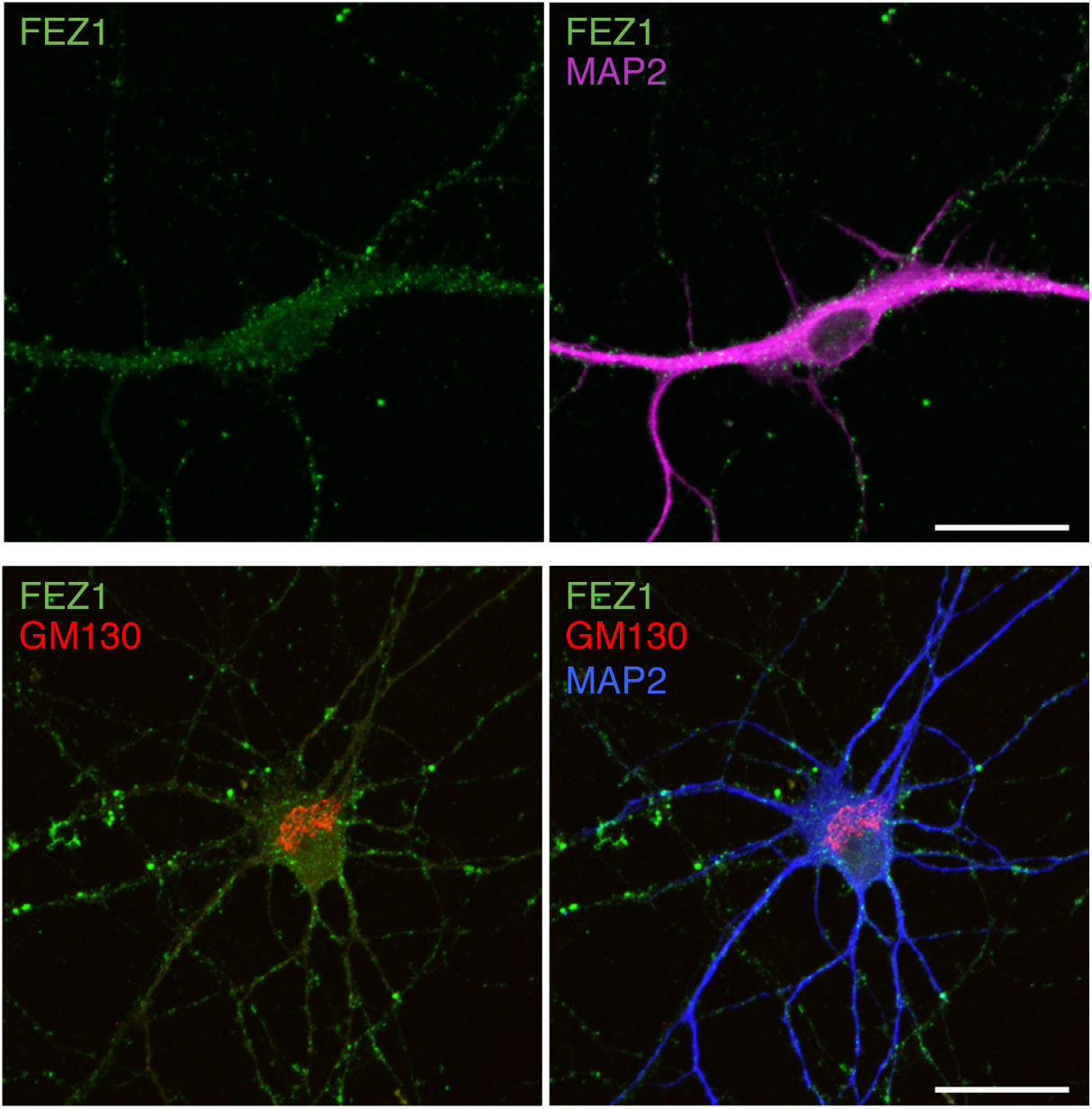
FEZ1 is expressed in the somatodendritic compartment of mammalian neurons. Images of DIV7 rat hippocampal neurons labeled with anti-FEZ1, anti-MAP2 to mark dendrites, and anti-GM130 to visualize Golgi apparatus; scale bars 35 μm.

